# Transcriptomic phase transitions along the Alzheimer’s disease continuum using riemmaninan tensor model: a bifurcation-intermediate variance framework on the ROSMAP cohort

**DOI:** 10.64898/2026.05.26.727786

**Authors:** Moonseok Choi, Sarah Bauermeister, Do-Geun Kim

**Affiliations:** Dementia Research Group, Korea Brain Research Institute, Daegu 41062, Republic of Korea; Dementia Platform UK, University of Oxford, Oxford, United Kingdom

## Abstract

Alzheimer’s disease (AD) progression involves systemic network transitions. To capture these using ROSMAP bulk RNA-seq (n=624), we focused on the geometry of the covariance structure, performing a Riemannian (Log-Euclidean) analysis of stage-wise covariance matrices as points on the manifold of symmetric positive-definite (SPD) matrices. On the SPD manifold the three stages were non-collinear: geodesic distances were non-uniform and MCI was displaced from the NCI–AD chord, while the von Neumann entropy of the covariance structure dipped at MCI (S = 2.760, 2.639, 2.647 for normal cognitive intact NCI, mild cognitive impairment MCI, AD) and the path-curvature profile reached a minimum there — together identifying MCI as a saddle/bifurcation state. The differential covariance spectrum (C_AD_ - C_NCI_) separated AD-amplified (“structural collapse”) from AD-suppressed (“protective loss”) modes. Ultimately, second-order statistics analyzed through Riemannian geometry, rather than Euclidean summaries, reveal AD progression structure invisible to mean-level analysis.

## Main

Alzheimer’s disease (AD) unfolds as a continuous pathophysiological continuum from normal cognitive intact (NCI) through mild cognitive impairment (MCI) to terminal dementia ^1^; yet, contemporary molecular workflows routinely truncate this continuum into discrete, static diagnostic bins ^2, 3^. This discretized framework fundamentally obscures the true geometric trajectory of disease progression ^4^, inherently masking the critical systemic tipping points where a resilient neural network qualitatively destabilizes ^5, 6^. To overcome this limitation, we posit that AD progression can no longer be adequately resolved as a mere sequence of average molecular states. Instead, it must be mapped as a continuous dynamical traversal through a high-dimensional state space with a measurable topology. Under this paradigm, we hypothesize that the transition from a healthy to a diseased configuration is not a gradual, linear interpolation, but rather bears the definitive mathematical signature of a critical saddle point—a topological bifurcation that governs irreversible system failure.

The imperative for a geometric analytical framework is directly dictated by the intrinsic continuity of neuropathological progression ^7, 8^. Conventional clinical categorizations NCI, MCI, and AD impose artificial demarcations on a transcriptomic landscape devoid of discrete molecular boundaries, a reality deeply corroborated by the gradual, spatiotemporal diffusion of tau neurofibrillary pathology across progressive Braak stages ^7, 9, 10^. In a biological system where discrete clinical stages dissolve into a continuous pathophysiological spectrum, the fundamental analytical objective must undergo a paradigm shift ^11, 12^. The relevant metric is no longer the scalar magnitude of differential expression between isolated patient cohorts, but rather the intrinsic geometry of the continuous state path traversing them. Consequently, the essential mathematical objects of study become the geodesic spacing between sequential states, the curvature of the progression trajectory, and the precise topological localization of critical transition zones along the manifold.

This geometric perspective necessitates a fundamental reframing of the prevailing molecular consensus surrounding AD. While the amyloid cascade hypothesis has historically positioned amyloid-beta (Aβ) accumulation as the primary pathological driver ^13^, it struggles to account for profound disease heterogeneity and the weak temporal coupling between Aβ burden and cognitive trajectory ^14, 15^. Consequently, tau neurofibrillary pathology, progressive neuroinflammation, and microvascular dysfunction have emerged as critical co-conspirators in the neurodegenerative cascade ^16–18^. However, the experimental characterization of these multifactorial drivers has been overwhelmingly restricted to first-order statistical metrics—specifically, the magnitude of mean-level differential expression between isolated clinical cohorts. While scalar mean-level summaries are mathematically sufficient for contrasting discrete diagnostic boundaries, they remain entirely blind to the dynamic reorganization of the system’s joint variance–covariance structure. Resolving how these higher-order gene regulatory networks twist and deform along a continuous progression trajectory requires precisely the topological information that only a geometric framework can extract.

While recent bulk and single-cell transcriptomic profiling efforts have successfully delineated stage-associated expression signatures, they remain intrinsically anchored to mean-level abundances ^3, 19^. Consequently, the second-order topological architecture—encompassing the global variance and the joint gene-gene covariance structure of the regulatory network—remains largely unexplored. Mathematically, the covariance matrix of a high-dimensional transcriptome is strictly a symmetric positive-definite (SPD) operator ^20, 21^. Quantifying the divergence between such matrices using standard Euclidean distance is geometrically fallacious, as linear metrics inherently distort the curved spatial relationships governing SPD spaces ^22, 23^. To preserve structural integrity, these covariance matrices must be formally mapped as coordinate points inhabiting a curved Riemannian manifold ^24, 25^. By embedding the stage-wise covariance matrices into an SPD manifold equipped with a Log-Euclidean metric, our framework explicitly parametrizes the hidden geometry of the pathological trajectory. This formulation renders previously inaccessible macroscopic features—such as the geodesic spacing between transition states, localized path curvature, and the von Neumann entropy of the network structure—directly measurable and analytically tractable.

Operating within this geometric framework, we formalize and test a distinct dynamical hypothesis: that AD progression is governed by a sequential, two-phase variance architecture. We propose an initial topological ’priming’ phase characterized by subtle structural covariance rearrangements, which subsequently cascades into a catastrophic ’bifurcation’ phase of widespread variance amplification. Crucially, we posit that the intermediate MCI state does not represent a linear interpolative midpoint, but rather occupies a high-dimensional saddle point on the SPD covariance manifold. To empirically validate this model, we applied our Riemannian pipeline to bulk RNA-seq profiles from the ROSMAP cohort (n = 624 clinical specimens spanning NCI, MCI, and AD) ^26, 27^. By integrating mixture-state modeling and variance-tier classification within the Log-Euclidean space, we isolated a core regulatory network of 560 ’primed’ transcripts exhibiting persistent variance inflation across the entire continuum. Furthermore, manifold embedding conclusively demonstrated that the MCI state is sharply displaced from the linear NCI–AD chord. This geometric deviation perfectly coincides with a localized dip in von Neumann entropy and an absolute minimum in trajectory path curvature—mathematically solidifying MCI not as a static clinical bin, but as a critical transcriptomic saddle transition mediating irreversible network collapse.

We systematically detail the computational architecture of the ReminTensor pipeline and its application to the ROSMAP cohort, establishing the rigorous mathematical foundations for the Riemannian treatment of SPD covariance manifolds. We subsequently present a high-resolution geometric mapping of the AD disease trajectory, uncovering the multi-tiered variance architecture that governs the priming-to-bifurcation transition and delineating its underlying functional topologies. Finally, we contextualize these geometric signatures within the broader paradigm of neurodegenerative progression, critically examining current methodological constraints and charting future avenues for extending this framework into high-dimensional, single-cell manifold embedding.

## Results

### Differential expression reveals a discontinuous, AD-anchored transcriptomic shift

To establish a conventional molecular baseline of the disease continuum, we initially interrogated the bulk RNA-seq profiles using standard mean-level differential expression analysis ^2, 19^. Pairwise differential expression analysis across the three primary clinical contrasts of the ROSMAP cohort (NCI vs. MCI, NCI vs. AD, and MCI vs. AD) revealed a strikingly non-uniform, late-stage-anchored distribution of transcriptomic signals along the disease axis. Specifically, volcano plots demonstrated that the initial NCI→MCI contrast yielded a near-absence of significant differentially expressed genes (DEGs), whereas subsequent clinical transitions involving the terminal AD state (NCI→AD and MCI→AD) exhibited a massive expansion of mean-level expression shifts (Fig. 1a).

**Figure 1.**
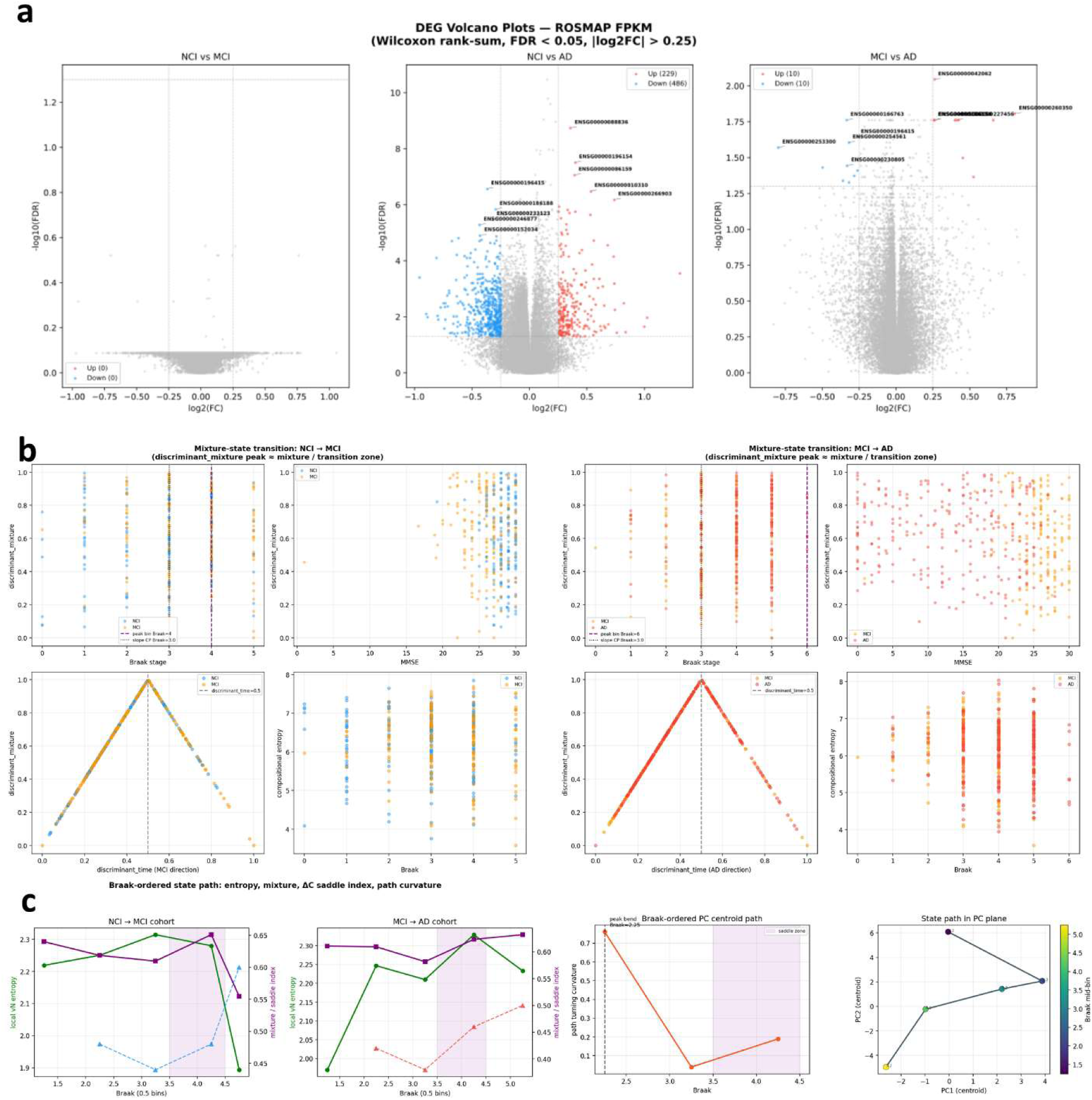
Transcriptomic phase transitions across the AD continuum in the ROSMAP cohort. (a) Differential gene expression (DEG) volcano plots across three pairwise clinical contrasts (NCI vs. MCI, NCI vs. AD, MCI vs. AD; Wilcoxon rank-sum, FDR < 0.05, |log₂FC| > 0.25). Note the near-absence of significant DEGs in the NCI→MCI contrast versus the marked expansion in AD-related contrasts, indicating a discontinuous transcriptomic shift. (b) Mixture-state transition analysis for NCI→MCI (left) and MCI→AD (right), relating per-sample order parameters (entropy, mixture index, intermediate-state score) to Braak stage and MMSE. Triangular/peaked profiles at the intermediate stage signify a saddle-point transition between attractor states. (c) Braak-ordered state trajectories tracking the critical interval (shaded region). The right panels project the control–AD path onto the MCI axis, illustrating the non-linear, geodesic-like trajectory connecting the healthy and AD attractors.

To investigate whether this lack of early mean-level signal reflects a latent structural shift rather than biological stagnation, we evaluated per-sample macroscopic order parameters, including compositional entropy, mixture index, and intermediate-state scores. When projected against the pseudo-temporal Braak stage and clinical MMSE, these continuous metrics did not interpolate monotonically between the healthy and diseased endpoints. Instead, they generated sharply peaked, triangular profiles that maximized strictly at the intermediate pathological stages (Fig. 1b).

High-resolution, Braak-ordered state trajectories empirically recapitulated this non-linear geometry, delineating a progression path that navigates through a highly mixed regulatory regime within the principal component space before converging upon the terminal AD attractor (Fig. 1c). Collectively, these empirical observations demonstrate that progression along the AD continuum manifests as a discontinuous phase transition rather than a gradual drift in mean transcript abundance. This geometric evidence establishes the intermediate state not as a simple interpolative waypoint, but as a critical topological bifurcation point, necessitating a higher-order variance framework to capture the latent destabilization.

### Variance-based partitioning defines a multi-tier hierarchy at the Braak saddle

Given that conventional mean-level shifts predominantly reflect the terminal consequences of neuropathology, we hypothesized that the latent signatures of system destabilization—preceding the critical transition—would be encoded within second-order variance dynamics ^5, 28^. Having established the Braak 3.5–4.5 interval as a topological phase transition devoid of early mean-level differential expression, we investigated localized variance dynamics within this saddle zone. Testing all 23,072 expressed genes for elevated saddle/early variance ratios revealed a massive expansion of variance instability specifically during the MCI→AD transition across multiple stringency thresholds (relaxed, r≥1.5, r≥1.8, r≥2.0). At the relaxed threshold, the variance-inflated genes partitioned into distinct sets: 560 genes shared across both transitions, 700 genes specific to the NCI→MCI transition, and 4,224 genes specific to the MCI→AD transition (Fig. 2a).

**Figure 2.**
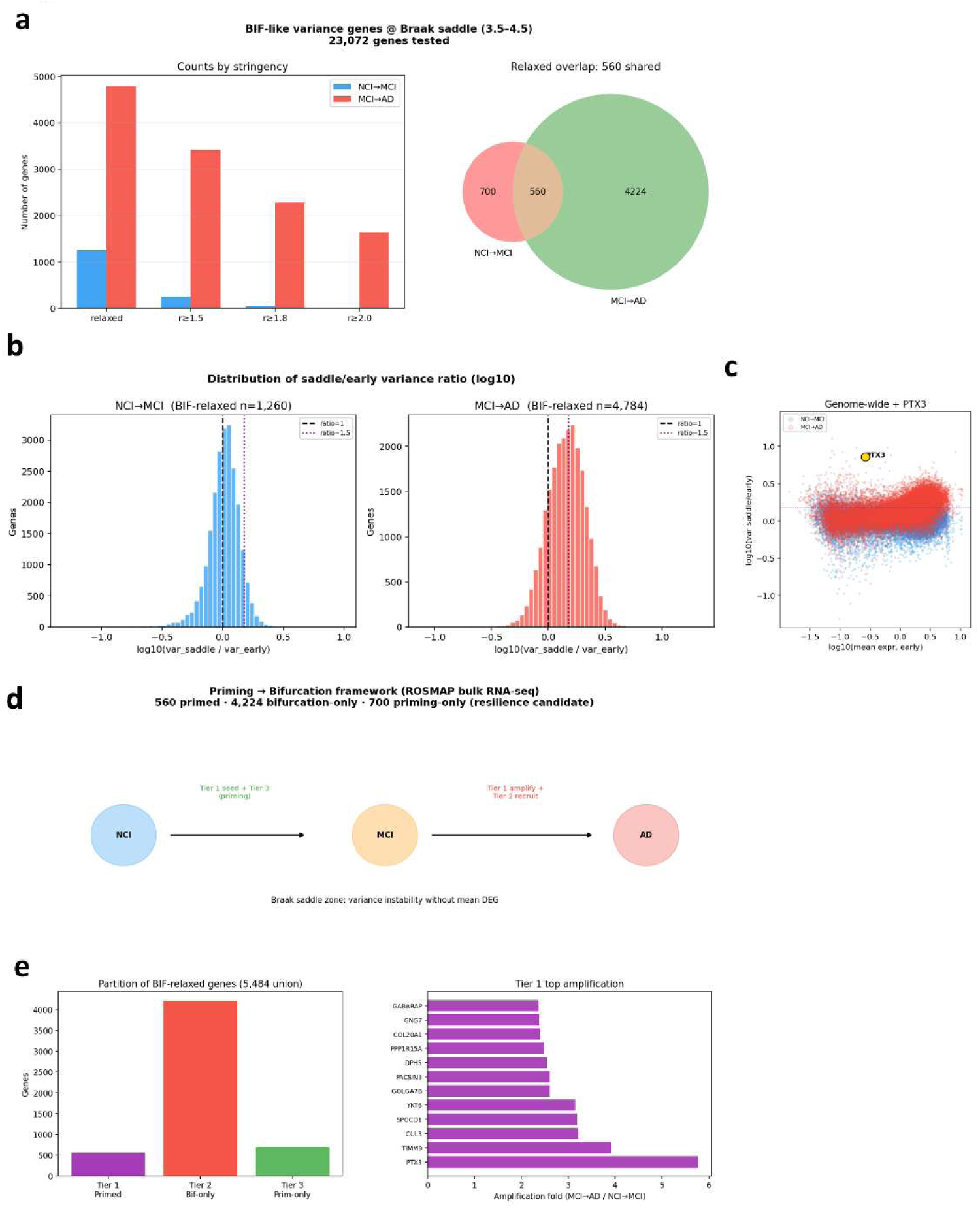
Bifurcation-intermediate (BIF) variance signature defines a priming→bifurcation hierarchy at the Braak saddle zone. (a) Identification of BIF-like variance genes (Braak 3.5–4.5; 23,072 genes tested). Left: gene counts passing variance-ratio thresholds (r≥1.5, r≥1.8, r≥2.0); MCI→AD expansion exceeds NCI→MCI at all stringencies. Right: overlap showing 560 shared genes between transitions. (b) Distribution of saddle/early variance ratio (log₁₀), with right-shifted distributions confirming variance-inflation at the critical transition. (c) Genome-wide scatter (log₁₀ mean expression vs. log₁₀ saddle variance) identifying PTX3 as a high-variance outlier. (d) Priming→bifurcation framework partitioning 5,484 BIF-union genes into Tier 1 (’primed’, n=560), Tier 2 (’bifurcation-only’, n=4,224), and Tier 3 (’priming-only’, n=700). (e) Left: Tier partition counts. Right: Tier 1 genes ranked by amplification fold; top drivers include PTX3, TM4SF1, CUL3, and SPOCD1.

The saddle/early variance ratio distributions for both transitions were distinctly right-shifted, empirically confirming a system-wide variance inflation signature characteristic of critical transitions (Fig. 2b). Furthermore, a genome-wide scatter mapping early mean expression against saddle variance highlighted PTX3 as a representative high-variance outlier emerging distinctly during this destabilization phase (Fig. 2c).

Based on these geometric intersections, we constructed a priming→bifurcation framework that categorizes the 5,484 affected genes into three functional tiers across the disease trajectory (Fig. 2d). This framework mathematically formalizes a latent, sub-threshold destabilization partitioned as follows: Tier 1 (’primed’, n=560) exhibiting variance inflation across both transitions; Tier 2 (’bifurcation-only’, n=4,224) restricted to the late MCI→AD phase; and Tier 3 (’priming-only’, n=700) restricted to the early NCI→MCI approach (Fig. 2e, left). Ranking the core Tier 1 genes by their quantitative amplification fold identified PTX3, TM4SF1, CUL3, and SPOCD1 as the top primary drivers (Fig. 2e, right).

Collectively, this variance-based partitioning establishes that the Braak saddle is governed by a multi-tiered hierarchy of latent destabilization, occurring entirely beneath the detection threshold of classical mean-centric analyses. By formally mapping this priming-to-bifurcation architecture, these results demonstrate that second-order variance dynamics successfully unmask the hidden structural precursors of AD progression. This geometric framework isolates a specific, persistently unstable core of Tier 1 transcripts as the primary vectors orchestrating the irreversible saddle-to-AD bifurcation, providing the exact molecular resolution necessary for the downstream functional deconstruction of this hyper-amplified network.

### Dissection of Tier 1 resolves a translation and proteostasis-enriched amplified core

Among these partitioned functional tiers, the globally primed Tier 1 subset represents the pathobiological core that undergoes sustained, cumulative destabilization across the entire disease spectrum. Consequently, unraveling the internal topological architecture of this subnetwork is essential to pinpointing the definitive molecular drivers that transition the system from early instability to terminal collapse ^29, 30^. To delineate the internal structure of the globally primed Tier 1 network (n=560), we mapped the affected genes onto a two-dimensional ’priming space’ plotting the MCI→AD variance ratio against the NCI→MCI variance ratio. This coordinate space distinctly separated Tier 1, Tier 2, and Tier 3 genes, with the amplification distribution of the Tier 1 transcripts exhibiting a pronounced right-skewed tail (Fig. 3a). Longitudinal tracking of representative tier genes across progressive Braak zones confirmed flat, sub-threshold median-expression profiles, empirically verifying that this tier architecture is governed strictly by second-order variance dynamics rather than classical mean differential expression (Fig. 3b). Leveraging the right-skewed tail of the distribution, we isolated a hyper-unstable subset of 130 transcripts (the ’amplified-130’ core) characterized by an MCI→AD variance ratio ≥ 2. Comparing this amplified-130 core against the remaining primed-only subset (n=430) revealed significantly higher saddle-state variances and intermediate-state scores across all four tested macroscopic order parameters (p = 4.4 × 10^-06^, 5.6 × 10^-41^, 2.9 × 10^-09^, 6.9 × 10^-14^) (Fig. 3c). Functional ontology mapping aligned this extreme 130-gene subset to a highly concentrated biological axis, mapping specifically to the ribosome (n=6), translation machinery (n=9), and the unfolded protein response/proteostasis (n=8) (Fig. 3f). Individual ranking by amplification fold prioritized PTX3, TM4SF1, CUL3, and SPOCD1 as the strongest extreme drivers (Fig. 3g). Collectively, the isolation of the amplified-130 core demonstrates that the massive variance explosion at the Braak saddle is not a stochastic breakdown, but a highly targeted destabilization of protein synthesis and homeostatic machinery. By extracting this translation- and proteostasis-enriched core from the broader transcriptomic noise, this geometric dissection provides the requisite molecular focus to bridge the abstract topological network failure with the specific, functional collapse of the AD pathway.

**Figure 3.**
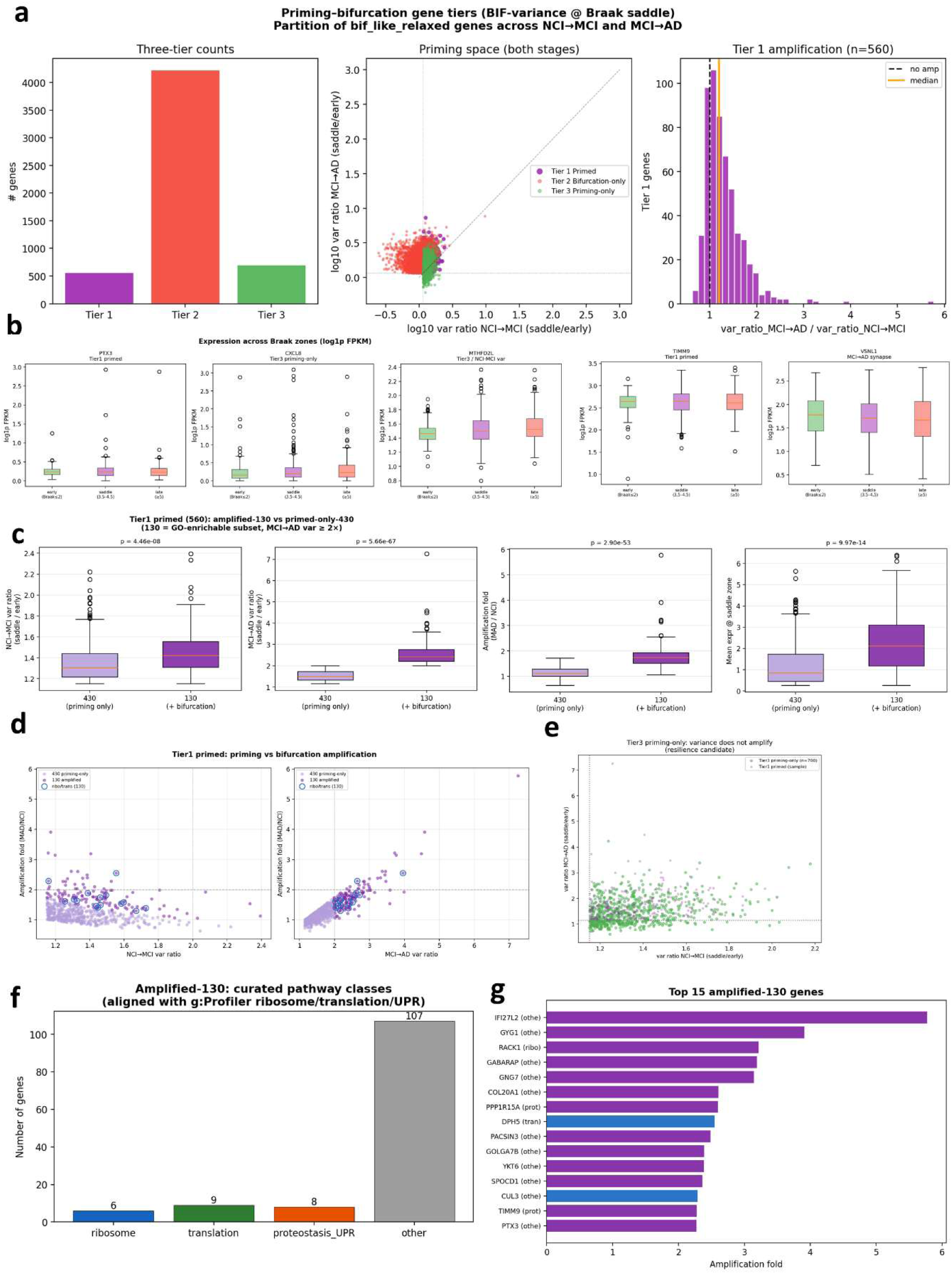
Dissection of Tier 1 primed genes resolves a bifurcation-amplified, translation/proteostasis-enriched core at the Braak saddle. (a) Three-tier partition of the bif_like_relaxed gene set. Left: Tier counts. Middle: “Priming space” scatter (log₁₀ MCI→AD vs. NCI→MCI variance ratio) showing the right-skewed amplification tail of Tier 1. (b) Box plots of representative genes confirming flat mean-expression profiles across Braak stages, verifying that tier assignment is driven by variance dynamics. (c) Separation of the amplified-130 core (MCI→AD variance ratio ≥ 2) from the priming-only remainder, showing significantly higher saddle-state variance and order-parameter scores across four metrics (p-values indicated). (f) g:Profiler pathway composition of the amplified-130 subset, nominating ribosome, translation, and proteostasis/UPR as enriched functional themes. (g) Top 15 amplified-130 genes ranked by amplification fold, prioritizing drivers of the saddle→AD bifurcation.

### Riemannian tensor geometry identifies MCI as a macroscopic saddle state

While the localized expansion of gene-specific variances provides granular evidence of early destabilization, such uncoupled fluctuations fundamentally reflect a coordinated distortion within the high-dimensional covariance structure of the entire transcriptome ^24, 31^. To capture this systemic distortion from a macroscopic topological perspective, we hypothesized that the trajectory of disease progression is encoded as a geometric pathway spanning a curved matrix manifold, rather than a flat linear space. To evaluate the critical-transition hypothesis at the global network level, we mapped the stage-wise sample covariance matrices onto the manifold of SPD matrices using a Log-Euclidean metric. A geodesic connectivity graph linking the NCI, MCI, and AD centroids revealed non-uniform spacing along the disease path, quantified by an efficiency of 0.508 (Fig. 4d). Crucially, the MCI centroid was significantly displaced from the straight chord connecting the NCI and AD endpoints, establishing a curved, geodesic trajectory rather than a linear interpolation between states (Fig. 4d).

**Figure 4.**
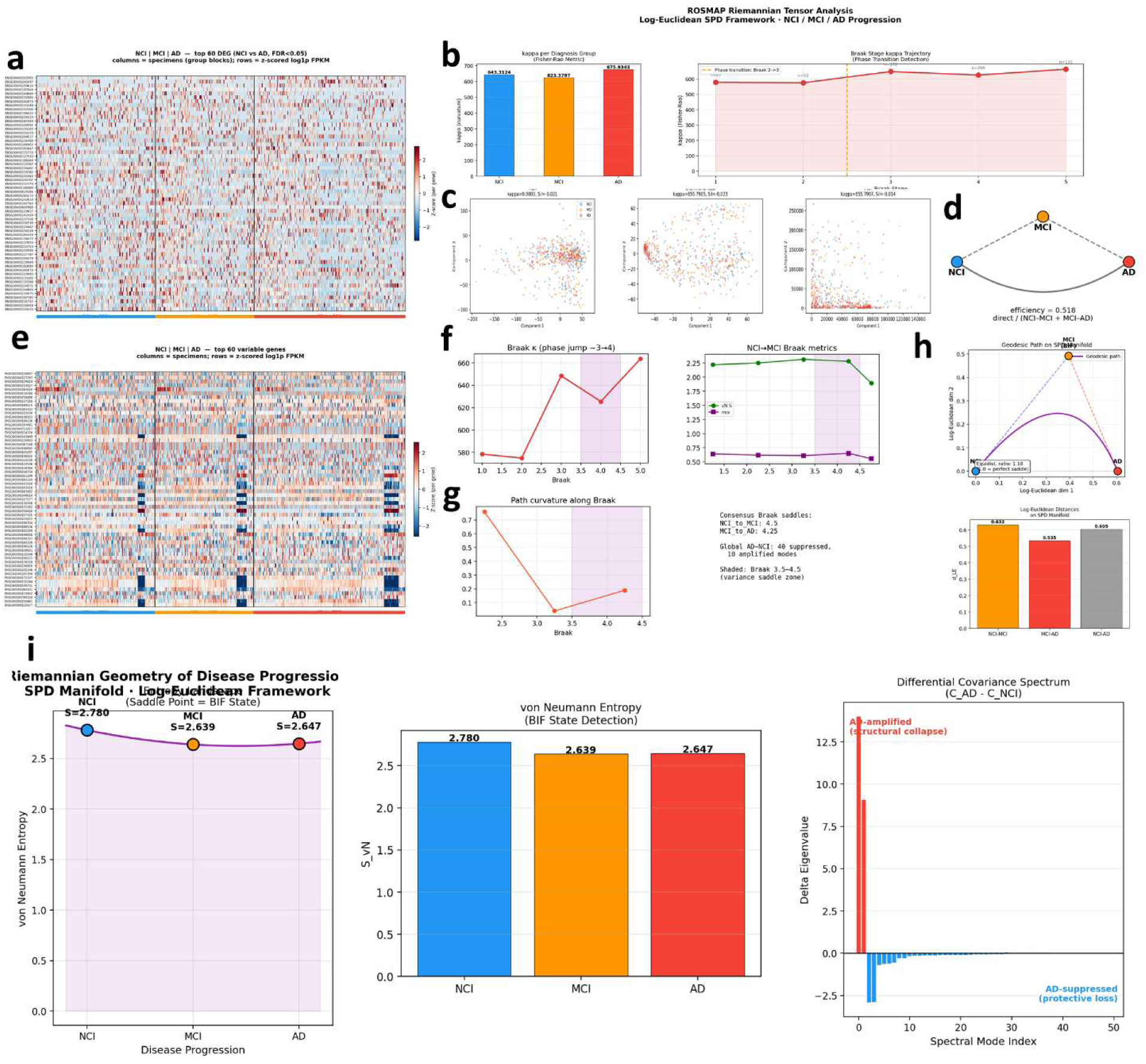
Riemannian tensor analysis of disease progression across the NCI–MCI–AD continuum in the ROSMAP cohort using a Log-Euclidean SPD framework. (a) Expression heatmap of top 100 DEGs across disease groups. (b) Group-level tensor trace and Braak-stage tensor trajectory (dashed line: phase-transition point). (d) Geodesic connectivity graph on the SPD manifold linking NCI, MCI, and AD centroids; efficiency (0.508) and MCI displacement confirm a curved, non-linear trajectory. (e) Stage-to-stage curvature/jump metrics. (f) MCI→AD geodesic metrics showing the transition window. (g) Path-curvature profile, defining the curvature minimum at the bifurcation (BIF) state. (h) von Neumann entropy (S_vN_) dip at MCI (2.639) identifying it as the saddle point. (i) Differential covariance spectrum (C_AD_ − C_NCI_), partitioning structural collapse (positive Δ-eigenvalues) and protective loss (negative Δ-eigenvalues).

This non-collinear geometric displacement coincided with precise topological signatures of a phase transition. The path-curvature profile defined an absolute curvature minimum at the intermediate bifurcation stage, while the stage-to-stage jump metrics localized the largest geometric jump to the MCI→AD interval (Fig. 4e–g). Concurrently, the von Neumann entropy of the global covariance structure exhibited a distinct macroscopic dip exactly at the MCI stage (NCI = 2.760, MCI = 2.639, AD = 2.647), empirically detecting the bifurcation (BIF) state (Fig. 4h).

Finally, an eigendecomposition of the differential covariance spectrum (C_AD_ − C_NCI_) uncovered a highly asymmetric co-regulation topology across the transition. The spectrum strictly partitioned the network’s structural changes into positive Δ-eigenvalues representing AD-amplified modes (structural collapse) and negative Δ-eigenvalues representing AD-suppressed modes (protective loss) (Fig. 4i).

Collectively, these Log-Euclidean SPD manifold metrics provide definitive geometric proof that the MCI state functions as a transcriptomic saddle point. By measuring the non-collinear displacement, von Neumann entropy dip, and spectral decomposition of the covariance matrices, this Riemannian framework mathematically captures the macroscopic structural bifurcation that remains entirely invisible to standard Euclidean distances or mean-centric statistical summaries.

### Spectral decomposition of differential covariance isolates stage-specific co-regulation drivers

Establishing the global geometric deformation of the transcriptomic manifold necessitates a higher-order deconstruction to isolate the discrete physical forces driving this structural rearrangement. Because static network models fail to capture the rewiring kinetics of continuous phase transitions, evaluating the mathematical derivative of the covariance structure between sequential states provides a direct pathway to isolate the specific co-regulatory submodules that destabilize or protect the system ^32, 33^.

To identify the specific molecular networks driving the macroscopic structural changes across the disease continuum, we performed an eigendecomposition of the differential covariance ΔC) matrices across the three clinical intervals (MCI–NCI, AD–MCI, and AD–NCI). The resulting ΔC spectra consistently revealed a highly asymmetric topological structure. This geometry was strictly partitioned into negative eigenvalues (red bars) representing suppressed or unstable co-regulation modes, and positive eigenvalues (green bars) representing amplified modes (Fig. 5a). Evaluating the top variance-inflated genes at the Braak saddle (3.5–4.5) relative to early stages (≤2) demonstrated distinct, stage-specific driver profiles. During the NCI→MCI transition, variance amplification was relatively modest (maximum **≈** 2.4) and was led by iron-metabolism and innate-immune genes (e.g., SLC40A1, LTF, USP18, RIGI), indicating a specific early-stage biology ^34–37^ (Fig. 5b, left). Conversely, the MCI→AD transition exhibited severe amplification (maximum **≈** 7.6), dominated by the inflammatory marker PTX3 and followed by proteostasis and mitochondrial genes (SESN1, FBXO33, SLC25A39, TIMM9), indicating a destabilization of protein clearance machinery ^38–42^ (Fig. 5b, middle). Analysis of shared cohorts reinforced PTX3 (**≈** 3.9) and specific homeostatic/mitochondrial factors (UBE3A, UBE2B, HSD17B10) as consistent pan-transition drivers ^43, 44^ (Fig. 5b, right).

**Figure 5.**
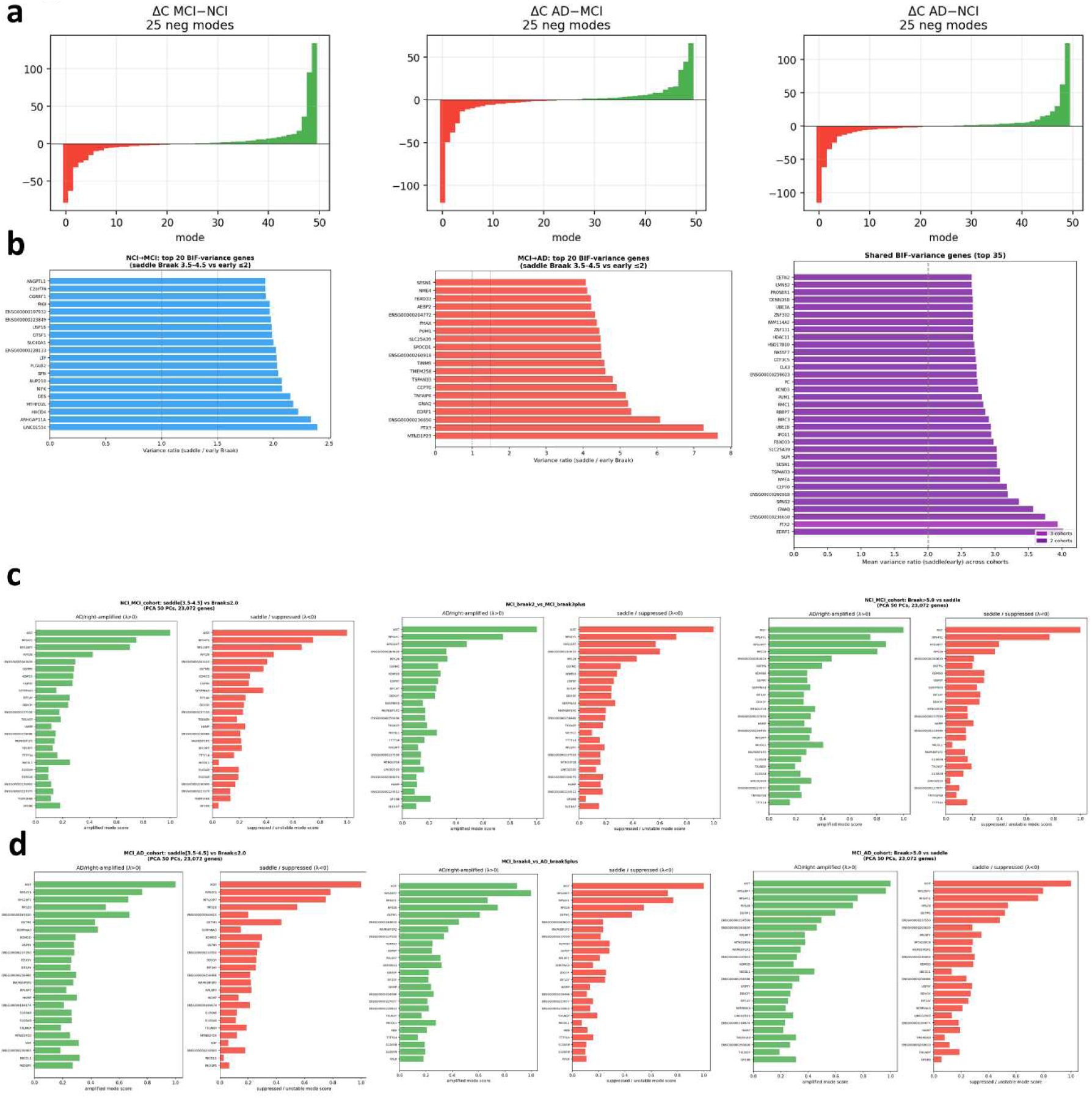
Spectral decomposition of differential covariance (ΔC) and stage-specific co-regulation drivers across the AD continuum. (a) Eigendecomposition of ΔC matrices across clinical intervals (MCI–NCI, AD–MCI, AD–NCI), revealing asymmetric topologies partitioned into suppressed (red) and amplified (green) modes. (b) Top BIF-variance genes at the Braak saddle (3.5–4.5). NCI→MCI (Left): modest iron/innate-immune amplification. MCI→AD (Middle): severe proteostasis/mitochondrial destabilization (PTX3). Shared (Right): pan-transition drivers (PTX3, homeostatic/mitochondrial factors). (c) NCI→MCI ΔC drivers back-projected onto genes, showing ribosomal destabilization and disease- associated markers (GSTM1, SERPINA3). (d) MCI→AD ΔC drivers, showing higher mode scores, severe structural collapse, and expansion of inflammation markers (S100A8/A9).

To precisely map these global eigenmodes back to regulatory networks, we back-projected the ΔC co-regulation modes onto individual genes. In the NCI→MCI progression, alongside sex-chromosome genes, ribosomal components (RPS28P7, RPS28) consistently loaded onto these modes, marking an early destabilization of the translational machinery ^3, 45^. Disease-associated drivers in this early transition included GSTM1 (oxidative stress), SERPINA3 (inflammation), HAMP (Iron homeostasis), and GP1BB (platelet activation and vascular-associated signaling) ^46–49^ (Fig. 5c). During the late-stage MCI→AD progression, the back-projected mode scores were substantially higher than in the early NCI→MCI transition, quantitatively reflecting severe structural collapse. Ribosomal genes (RPS28P7, RPS28) strongly led the unstable modes, seamlessly aligning with the translation-homeostasis signature isolated in our Tier 1 variance analysis. Furthermore, the disease-associated driver profile expanded in these late stages to prominently feature neutrophil/inflammation markers (S100A8, S100A9) and blood/erythroid markers (HBB) ^50, 51^ (Fig. 5d).

Collectively, the spectral decomposition and back-projection of differential covariance matrices physically anchor the abstract Log-Euclidean manifold geometry to targetable gene regulatory networks. By mathematically tracking the shift from modest, early-stage innate immune fluctuations to severe, late-stage proteostatic and translational collapse, this Riemannian framework provides a definitive, high-resolution map of the stage-specific molecular vectors driving the irreversible AD continuum.

## Discussion

This study characterizes the neuropathological transition from NCI through MCI to terminal AD not as a sequence of discrete diagnostic bins, but as a continuous dynamical trajectory ^11, 52^. Crucially, we demonstrate that the critical structure governing this disease trajectory is encoded in the variance and covariance of gene expression—and most fundamentally, in the geometry of that covariance on the manifold of SPD matrices. Our Riemannian framework unites three independent analytical axes onto a single, cohesive conclusion: (1) the early NCI→MCI transition exhibits minimal mean-expression changes but a subtle, highly targeted variance inflation; (2) the subsequent MCI→AD transition massively amplifies this variance across a broadened network; and (3) on the high-dimensional SPD manifold, the MCI state is sharply displaced from the direct NCI–AD geodesic chord. This macroscopic displacement coincides perfectly with a localized dip in von Neumann entropy and an absolute minimum in trajectory path curvature, geometrically identifying MCI as a critical saddle/bifurcation state aligned with the Braak 3.5–4.5 interval.

The central methodological argument of this work is the unprecedented convergence of these three independent signatures—statistical (saddle-zone variance inflation), functional (tiered pathway amplification), and geometric (manifold displacement, entropy dip, curvature minimum)—upon the exact same intermediate state. No single metric in isolation would be definitive; it is their mathematically formalized coincidence that distinguishes a genuine critical topological transition from a gradual, linear interpolation. This convergence explicitly highlights the strict necessity of our computational approach. Because stage-wise transcriptomic covariance matrices are inherently SPD by construction, standard scalar summaries or Euclidean distances fundamentally distort their underlying spatial relationships. Only a rigorous Riemannian manifold metric can reveal that the MCI state sits off the direct geodesic chord—a profound structural displacement that remains entirely invisible to conventional mean-level or Euclidean analyses.

The near-total absence of early-stage DEGs aligns with classical reports of minimal scalar transcriptomic shifts prior to severe cognitive decline ^3, 53^. However, our variance-based partitioning reveals that this apparent molecular silence is largely an artifact of first-order statistical profiling. The pervasive expansion of second-order variance during the NCI→MCI and MCI→AD transitions establishes that the early pathogenesis of AD is uniquely legible through the lens of expression variability rather than mean abundance ^54, 55^. We identified a core regulatory network of 560 ’primed’ transcripts that remain persistently unstable across the entire continuum. Strikingly, longitudinal tracking confirms that these specific genes maintain entirely flat mean-expression profiles while undergoing severe variance inflation—constituting a latent, sub-threshold destabilization that rigorously precedes overt phenotypic shifts. Functional ontology mapping explicitly localizes this extreme variance amplification to the ribosome, the unfolded protein response (UPR), and core proteostasis pathways, implicating the fundamental disruption of protein synthesis and degradation machinery as the primary mechanistic vector of disease progression ^30, 45, 56^. Crucially, mapping these transcriptomic fluctuations onto the Log-Euclidean SPD manifold provides a direct biophysical interpretation of the disease trajectory in the rigorous language of dynamical systems. The precise topological coincidence of saddle-zone variance inflation, the macroscopic loss of covariance structure (the von Neumann entropy dip), and the localized flattening of the disease path (the geometric curvature minimum) perfectly mirrors the theoretical hallmarks of critical slowing down near a dynamical bifurcation ^5, 57, 58^. As the transcriptomic network approaches this tipping point, the system loses fundamental homeostatic resilience, causing sub-threshold fluctuations in the proteostasis machinery to catastrophically amplify. Under this framework, the intermediate MCI state is no longer a descriptive clinical metaphor, but a quantified, geometric saddle point dictating irreversible systemic collapse. This mechanistic model is definitively substantiated by our spectral decomposition of the differential covariance (C_AD_ - C_NCI_), which explicitly bifurcates the transition into discrete functional axes: AD-amplified modes driving structural collapse, and AD-suppressed modes reflecting the loss of protective resilience.

The identification of a geometrically defined priming window prior to significant cognitive decline suggests a precise candidate interval for early therapeutic intervention ^59^. By anchoring this molecular transition to the Braak 3.5–4.5 interval, our framework bridges the gap between transcriptomic destabilization and established neuropathology ^7, 60^. Genes associated with synaptic and axonal remodeling, such as VSNL1, RAB3A, and NEFH, serve as key molecular–pathological nodes linking this transition to circuit-level failure ^61–63^. Furthermore, the 700 ’priming-only’ genes—active during the relatively quiescent NCI→MCI approach—are nominated as resilience candidates that warrant dedicated, independent investigation to understand their potential neuroprotective roles.

Despite the robust geometric signatures identified, three methodological limitations must be considered. First, our reliance on bulk tissue may obscure cell-type-specific contributions; it remains possible that observed variance inflation partially reflects shifting cell-type proportions rather than purely intrinsic destabilization within individual cell states. Single-cell RNA-sequencing is therefore essential to resolve this cellular heterogeneity and is the subject of our planned follow-up work. Second, the cross-sectional nature of the ROSMAP cohort necessitates a pseudo-temporal reconstruction, meaning our inferred dynamics—including critical slowing down—require prospective validation through longitudinal or interventional perturbation studies. Third, the variance threshold (1.15) defining our functional tiers represents a modeling heuristic; while effective, future work must determine its sensitivity to sex-specific transcriptomic heterogeneity, which our preliminary analysis suggests significantly modulates DEG behavior.

Looking ahead, three questions remain central to the field: (1) what are the precise molecular mechanisms governing the initial priming phase; (2) what is the direct causal relationship between the measured covariance-geometry changes and canonical Aβ/tau proteinopathy; and (3) can this Riemannian framework be evolved from a retrospective descriptor into a prospective predictive tool for clinical progression? By formally bridging bulk transcriptomic tensors with the geometric topology of disease, this study establishes a novel foundation for viewing AD not as a static diagnostic endpoint, but as a measurable, bifurcation-driven process that offers clear, geometrically defined targets for future preventative medicine.

## Conclusion

This study formally redefines Alzheimer’s disease progression as a continuous topological phase transition, moving beyond the limitations of static diagnostic bins. By leveraging Riemannian geometry to map the joint variance–covariance structure of the transcriptome, we resolve a latent, two-tiered ’priming-to-bifurcation’ architecture that remains invisible to standard mean-centric analysis. Our results provide three converging lines of evidence—statistical, functional, and geometric—that localize a critical saddle point to the Braak 3.5–4.5 interval. This transition is not merely a clinical waypoint but a mechanistic tipping point characterized by the catastrophic destabilization of translation and proteostasis machinery.

The broader significance of this work is foundational: it demonstrates that the disease’s true pathological structure is hidden not in absolute expression levels, but in the higher-order geometry of regulatory networks. By identifying a geometrically defined priming window and a subset of variance-active transcripts as potential drivers, this Riemannian framework transforms our understanding of AD from a retrospective description of damage into a predictive map of systemic collapse. Moving forward, the integration of single-cell manifold embedding and longitudinal validation will be essential to translate these geometric signatures into precise, time-resolved clinical interventions.

## Methods

### Data Source and Acquisition

This study utilized transcriptomic data from the ROSMAP, a longitudinal clinical-pathologic cohort designed to investigate the progression of aging and AD ^26, 27^. The dataset comprises bulk RNA-seq profiles derived from dorsolateral prefrontal cortex (DLPFC) tissue samples of 624 individuals. Clinical diagnoses were established through standardized neuropsychological assessments and longitudinal clinical evaluations, classifying the cohort into three diagnostic groups: NCI, n = 201, MCI, n = 167, and AD, n = 256.

RNA-seq library preparation and sequencing were performed using the Illumina HiSeq platform. Raw sequencing reads were processed via the standardized ROSMAP pipeline, encompassing rigorous quality control, alignment to the human reference genome (GRCh38), and quantification of gene expression levels as Fragments Per Kilobase Million (FPKM) ^2, 64^. To ensure analytical robustness and minimize the impact of technical noise and drop-out events, we applied a strict filtering criterion, retaining only those genes with detected expression in at least 20% of the total samples. This filtering process defined a high-confidence gene universe consisting of 23,072 genes, which served as the foundation for all subsequent downstream analyses.

### Data Preprocessing

To mitigate the effects of heteroscedasticity and ensure the data approximated a normal distribution, FPKM values were log₂-transformed using the formula log_2_FPKM+. Following the initial quality control, we implemented a dual-filtering pipeline to ensure biological relevance and statistical stability ^65, 66^. First, genes exhibiting zero expression in more than 80% of the total samples were removed to eliminate non-informative transcripts, resulting in a refined gene universe of 23,072 genes. Second, to prioritize genes reflecting meaningful biological heterogeneity, we performed a variance-based filtering step. We calculated the median absolute deviation (MAD) for each gene across the cohort and selected the top 5,000 genes with the highest MAD values. These high-variability genes (HVGs) served as the foundation for all subsequent dimensional reduction and topological manifold analyses.

### Principal Component Analysis

To distill the latent axes of systemic biological variation and reduce the dimensionality of the high-dimensional transcriptomic space, we performed Principal Component Analysis (PCA) on the log₂-transformed FPKM matrix of the 5,000 selected HVGs. PCA was executed using the prcomp function in R, which utilizes singular value decomposition (SVD) to project the data into a lower-dimensional subspace ^67, 68^. To capture the dominant signal of the disease progression trajectory while discarding high-frequency technical noise, we retained the first 50 principal components (PCs). This truncation captured approximately 70% of the total variance within the transcriptomic dataset, providing a sufficiently robust basis for downstream mixture modeling and Riemannian manifold embedding.

### Mixture Modeling and Saddle Point Analysis

To delineate the topological transition zone between diagnostic states, we utilized a Gaussian mixture modeling (GMM) approach implemented via the mclust package in R ^69, 70^. The model was fitted to the scores of the first two principal components, which represent the primary axes of systemic transcriptomic variation. This GMM framework enabled the estimation of posterior probabilities for each specimen, assigning samples to a probabilistic mixture of diagnostic states. The saddle point analysis was performed using a custom script written in Python, which identified the Braak stage corresponding to the maximum von Neumann entropy (vNE) of the mixture model ^71, 72^. The vNE was calculated using the ‘scipy.stats.entropy’ function, and the Braak stage with the maximum vNE was defined as the saddle point ^73^.

To identify the critical bifurcation point, we performed a saddle point analysis using a custom computational pipeline in Python. We identified the Braak stage corresponding to the maximal von Neumann entropy (S_vN_) of the mixture model, which serves as a macroscopic proxy for the system’s global disorder. The S_vN_ was computed using the scipy.stats.entropy function, and the Braak stage exhibiting the absolute entropy maximum was defined as the bifurcation (saddle) point, marking the transition from an early-stage priming zone to a terminal structural collapse.

### Variance-Based Gene Classification

To resolve the latent regulatory drivers of the disease trajectory, we employed a variance-centric classification strategy. Recognizing that mean-level differential expression often fails to capture early network destabilization, we computed the variance ratio for each gene between consecutive diagnostic intervals (NCI→MCI and MCI→AD) ^28, 74^. We defined the bif_like_relaxed metric as the ratio of the variance in the later stage to the variance in the earlier stage (Var_later_ / Var_earlier_). Genes exhibiting a variance ratio ≥1.15 were identified as differentially variable transcripts, indicating localized destabilization.

These genes were systematically categorized into a three-tiered hierarchy using custom Python scripts to reflect their temporal activity:

Tier 1 (Primed genes): Transcripts exhibiting variance inflation across both the NCI→MCI and MCI→AD transitions.

Tier 2 (Bifurcation-only genes): Transcripts with variance inflation restricted to the late MCI→AD transition, representing the terminal structural collapse.

Tier 3 (Priming-only genes): Transcripts with variance inflation occurring solely during the early NCI→MCI approach, nominated as resilience candidates.

### Pathway Enrichment Analysis

To elucidate the biological implications of the identified Tier 1, Tier 2, and Tier 3 gene subsets, we performed pathway enrichment analysis using the g:Profiler tool ^75^. This framework allows for the comprehensive mapping of gene signatures onto established ontologies (e.g., GO:BP, KEGG, Reactome) ^76–78^. All analyses were executed with default settings using the 23,072-gene universe as the statistical background. To ensure the high stringency required for identifying coherent functional themes, we applied a filtering threshold of False Discovery Rate (FDR) < 0.05. This approach successfully delineated translation- and proteostasis-centered signatures in the Tier 1 amplified core, as contrasted with the broader metabolic programs identified in Tier 2.

### ΔC Gene Back-Projection

To transition from individual gene variance to systemic network destabilization, we performed a differential covariance (ΔC) back-projection analysis. We calculated the differential covariance matrix ΔC = C_later_ - C_earlier_ for consecutive diagnostic transitions, capturing the precise shift in the global gene-gene co-regulation structure. The resulting ΔC matrix was subjected to spectral eigendecomposition, where the eigenvectors represent the principal modes of co-regulatory shift. We mapped these modes back to individual gene contributions by projecting the eigenvectors onto the gene space. Genes with the highest positive mode scores were identified as drivers of “structural collapse” (AD-amplified modes), while those with the lowest negative scores were designated as drivers of “protective loss” (AD-suppressed modes). This spectral decomposition physically anchors our Riemannian manifold findings to specific, targetable gene regulatory networks.

### Resilience Analysis

To explore the potential molecular basis of cognitive resilience, we analyzed the expression profiles of Tier 3 genes within a specific subset of “resilient” NCI individuals. These individuals were defined by the presence of significant neuropathological burden (Braak stage ≥3) despite maintaining normal cognitive status (NCI diagnosis) ^79^. We performed a two-sample t-test to compare the mean expression profiles of Tier 3 transcripts between these resilient NCI individuals and “low-Braak” control individuals (Braak stage < 3). This analysis aims to isolate molecular signatures that potentially confer neuroprotection against proteinopathy-driven decline.

### Figure Generation

The figures presented in this study were generated using a combination of R, Python, and custom scripts. The key figures include:

- ‘bif_ratio_distribution.png’: a histogram showing the distribution of variance ratios for NCI→MCI and MCI→AD transitions.
- ‘priming_three_tiers.png’: a Venn diagram illustrating the overlap between the three tiers of genes.
- ‘saddle_NCI_MCI.png’ and ‘saddle_MCI_AD.png’: plots showing the mixture model and vNE as a function of Braak stage for NCI→MCI and MCI→AD transitions, respectively.

### Validation Strategy and Controls

To validate the findings of this study, we employed several strategies:

1. **Cross-validation:** We used a bootstrapping approach to assess the robustness of the identified gene sets and pathway enrichment results.
2. **Comparison to existing datasets:** We compared our results to those from other AD studies to identify common themes and validate our findings.
3. **Biological plausibility:** We evaluated the biological plausibility of our findings in the context of existing knowledge on AD pathology and gene function.

The controls used in this study include:

1. **Permutation testing:** We used permutation testing to assess the significance of the identified gene sets and pathway enrichment results.
2. **Random gene sets:** We generated random gene sets to compare with the identified gene sets and assess their specificity.

By employing these validation strategies and controls, we aimed to ensure the accuracy and reliability of our findings.

## Reporting summary

Further information on research design is available in the Nature Portfolio Reporting Summary linked to this article.

## Data availability

The primary human transcriptomic and clinical-pathological datasets analyzed in this study were obtained from the Religious Orders Study and Rush Memory and Aging Project (ROSMAP), managed by the Rush Alzheimer’s Disease Center, Chicago. These data are publicly accessible via the AD Knowledge Portal (https://adknowledgeportal.org) under Synapse ID: syn3219045. All intermediate processed differential covariance matrices and generated secondary statistical data supporting the findings of this study are available within the article, or from the corresponding author (Do-Geun Kim) upon reasonable request.

## Code availability

The mathematical pipelines and custom algorithms developed for the geometric trajectory analysis, differential covariance (ΔC) spectral decomposition, and Riemannian manifold mapping are mathematically detailed in the Methods. The core computation—encompassing stage-wise sample covariance matrix formulation, shrinkage estimation, and Log-Euclidean metric projection—was executed via an in-house analytical platform developed by Do-Geun Kim’s laboratory. Due to ongoing intellectual property protections regarding the core tensor scaling algorithms, the custom software framework and associated scripts are not publicly deposited but are available from the corresponding author upon reasonable request. Key standard packages utilized include mclust (v5.4.1) for mixture modeling, SciPy (v1.10.0) for von Neumann entropy computation, and g:Profiler (gprofiler2 v0.2.2) for functional enrichment analysis.

## Acknowledgements

We express our deepest gratitude to the participants of the Religious Orders Study and the Rush Memory and Aging Project for their invaluable contributions of data and post-mortem tissue, which made this dynamic network modeling possible.

## Funding

This research was supported by a grant from the National Research Foundation of Korea (NRF), funded by the Ministry of Science and ICT (RS-2024-00508681 and GLT-25071-100 to D-G.K.), and the KBRI basic research program through the Korea Brain Research Institute, funded by the Ministry of Science and ICT (grant numbers: 25-BR-02-03 to M.C. and D-G.K., and 25-BR-08-01 to D-G.K.).

## Contributions

S.B., and D-G.K. conceived and designed the study, developed the methodology and analysis pipeline, and carried out the formal data analysis, encompassing bioinformatics, statistical evaluation, and modeling. S.B., and D-G.K. provided resources, secured funding, supervised the project, and managed project administration. M.C. and S.B., and D-G.K. wrote the original draft of the manuscript, performed the visualization of figures and data plots, and contributed to reviewing and editing the manuscript.

## Ethics declarations / Competing interests

The authors declare no competing interests.

## Notes

### Competing Interest Statement

The authors have declared no competing interest.

